# Multi-ancestry polygenic risk scores using phylogenetic regularization

**DOI:** 10.1101/2024.02.14.580313

**Authors:** Elliot Layne, Shadi Zabad, Yue Li, Mathieu Blanchette

## Abstract

Accurately predicting phenotype using genotype across diverse ancestry groups remains a significant challenge in human genetics. Many state-of-the-art polygenic risk score models are known to have difficulty generalizing to genetic ancestries that are not well represented in their training set. To address this issue, we present a novel machine learning method for fitting genetic effect sizes across multiple ancestry groups simultaneously, while leveraging prior knowledge of the evolutionary relationships among them. We introduce DendroPRS, a machine learning model where SNP effect sizes are allowed to evolve along the branches of the phylogenetic tree capturing the relationship among populations. DendroPRS outperforms existing approaches at two important genotype-to-phenotype prediction tasks: expression QTL analysis and polygenic risk scores. We also demonstrate that our method can be useful for multiancestry modelling, both by fitting population-specific effect sizes and by more accurately accounting for covariate effects across groups. We additionally find a subset of genes where there is strong evidence that an ancestry-specific approach improves eQTL modelling.

## Introduction

With the growing scale of large biobanks across heterogeneous human populations, there is an urgent need to develop polygenic risk score (PRS) and other genotype-to-phenotype prediction methods that can accurately capture the multiancestry genetic effects on diverse phenotypes, ranging from expression quantitative traits to complex diseases. However, existing PRS methods perform best only when trained and evaluated on populations with a homogeneous ancestry [Martin et al., 2019]. The amount of training data available outside of European and East Asian populations remains small. As a result, the majority PRS studies are performed using data exclusively from one of these groups, with European ancestry historically receiving particular attention in the literature. Consequently, popular PRS methods tend to have significantly poorer performance in populations that are underrepresented in large genomic databases, potentially contributing the health inequities [Duncan et al., 2019]. The challenge of cross-ancestry prediction has also been observed in the context of expression quantitative trait loci (eQTL) modelling [Mogil et al., 2018].

There are multiple mechanisms that may be contributing to this poor generalization across populations. These include changes in allele frequencies, bias in SNP selection during genome-wide association studies (GWAS) towards SNPs prevalent in large training populations, and changes in effect sizes due to differences in genetic architecture or interaction effects [Timpson et al., 2018, Kachuri et al., 2023]. Changes in linkage disequilibrium (LD) due to differences in ancestry composition between the training and test data can also lead to poor generalization [Guo et al., 2018, Wang et al., 2020]. In this work we present DendroPRS, a novel method for fitting SNP effect sizes across multiple ancestry groups. In DendroPRS, SNP effect sizes are allowed to evolve along the branches of the phylogenetic tree relating the different ancestry groups [Duda and Zrzavý, 2016], but subject to a novel tree-based regularization scheme. Tree-distance between ancestry groups can be used as a useful inductive bias, informing how much variance we may expect in effect sizes across different populations.

We first validate the ability of DendroPRS to fit shifting effect sizes across multiple populations in a simulation study. We then further explore our method’s performance on modelling gene expression across five different populations on Geuvadis data from the 1000 Genomes Project (1Kg) [Consortium et al., 2015, Stranger et al., 2012, Lappalainen et al., 2013], and find evidence that there exist a small number of genes whose expression is likely regulated in a population-specific manner. We further test DendroPRS on the task of fitting PRS models simultaneously across different sub-populations within the UK Biobank (UKB) [Bycroft et al., 2018], finding evidence that DendroPRS can outperform baseline methods on certain phenotypes when used to fit a limited number of highly significant SNPs, along with covariates. Finally, we release on implementation of DendroPRS for community use.

## Related Work

Multiple existing methods seek to address the challenge of cross-population polygenic prediction [Wang et al., 2022]. Most existing approaches to tackle this problem operate by inferring ancestry-specific effect sizes while imposing shrinkage rules or priors that encourage concordance in effects across ancestry groups. One of the most widely-used methods in this space is PRS-CSx [Ruan et al., 2022] a Bayesian method that imposes cross-ancestry continuous shrinkage prior with parameters fit using GWAS summary data across multiple populations. Another popular approach in this space is exemplified by the XPA and Vilma methods [Cai et al., 2021, Spence et al., 2022]. These methods model the effect of each variant via mixtures of multivariate Gaussians, with covariance between the populations determined using the cross-population correlation of the trait under consideration. The XP-BLUP method fits a multicomponent linear mixed model, in which SNPs are assigned to a component using pooled data from all population, and a best unbiased linear predictor (BLUP) is fit for each population independently, [Coram et al., 2017]. As a way to bypass the difficulties posed by spurious correlations due to LD, Polypred, introduced by Weissbrod et al. [2022], combines a fine-mapping based approach leveraging functional annotations with an LMM predictor.

Orthogonally, recent work explored transfer learning approaches, in which a PRS inferred from a large population is fine-tuned for a smaller target population. Zhao et al. [2022] present such an approach. They first fit a set of effect sizes for an auxiliary population with large sample size (e.g. European), and then update the effect sizes via stochastic gradient descent, using predictive performance in a second, target population as training signal.

We note that, with the exception of LMM-style methods, previous work for cross-population PRS predominantly considers the setting of a large auxiliary population and a single small target population. Our proposed method is unique in that we leverage knowledge of the historical relationships among different ancestry groups in order to better fit effect sizes simultaneously across all populations in the analysis. By accounting for the degree of genetic similarity across all sub-populations, we can improve predictive performance, in comparison to regimes that fit effect sizes for each target population independently.

The task of cross-ancestry eQTL has been less thoroughly explored than PRS. Mogil et al. [2018] confirm that the genetic architecture of gene expression traits is correlated across subpopulations with strength that is proportional to shared ancestry, motivating the use of our method. Additionally, Zeng et al. [2022] explore use of an LMM eQTL model applied to multiple cohorts as part of a fine-mapping pipeline integrating both GWAS and eQTL information.

In contrast to the above methods, a key novelty of our work is the use of tree-guided regularization to fit correlated genetic effect sizes across different populations. DendroPRS represents, to our knowledge, the first use of a phylogenetic regularization for this task. Conceptually similar regularization schemes were used by Layne et al. [2020] for improving the generalization of models trained on multi-species tasks such as inferring antibiotic resistance across bacterial species, and by Zitnik and Leskovec [2017] for learning molecular embeddings capturing function across hierarchies of tissues.

## Methods

### Tree-guided multi-ancestry effect sizes

We consider that we have access to a *n × m* genotype matrix **X** consisting of *n* individuals, each with *m* SNPs, and a vector **y** of *n* corresponding real-valued phenotypes.

We further assume there exists a set of populations *𝒫*, such that each individual *i ∈*{1, …, *n*} is assigned to some population *p ∈ 𝒫*. We denote **X**^(*p*)^ and **y**^(*p*)^ as the subset of rows of **X** and **y** corresponding to individuals belonging to population *p*.

We assume SNPs have additive effects on **y**, with population-specific genetic effect sizes β_*p*_ and noise term *ϵ*:

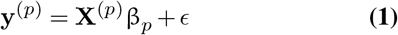

The populations *𝒫* consist of a set of ancestry groups. Human ancestry groups share a well-understood set of historical relationships, which can be reasonably depicted as a tree for non-admixed populations [Duda and Zrzavý, 2016]. We denote the binary tree 𝒯 = (𝒱, *ε*), where *𝒱* and *ε* are respectively the nodes and edges in *𝒱*. For each population *p ∈ 𝒫*, there exists a corresponding leaf node *v*_*p*_ *∈ 𝒱*, and the topology of *𝒯* reflects the historical relationships between all members of *𝒫*.

### Model Formulation

DendroPRS consists of a set of learnable parameters at the root of the tree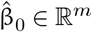, and a set of learnable “off-set” parameters 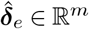 for each edge *e ∈ ε*. For each population *p*, we define *𝒮*_*p*_*⊂ ε* as the set of edges making up the path from the root of *𝒯* to the leaf node *v*_*p*_. The estimated genetic effect sizes for a population *p* are modeled as the sum of the root-level effect sizes 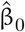 and their updates along the path *𝒮*_*p*_:

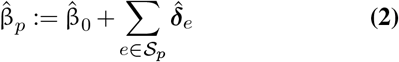

For each *p ∈𝒫*, we seek to fit a set of population-specific effect sizes 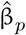 with sparsity-inducing regularization on each ***δ***, such that 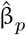 and 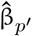 will be increasingly similar the closer *v*_*p*_ and 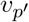 are located in *𝒯*. To this end, DendroPRS employs three regularization loss terms:

We assume that changes in effect sizes are likely to be sparse across edges in *𝒯*. Therefore, we impose an L1 penalty on the offset parameters at each edge.

1. As many SNPs may have an effect size of zero, we further apply a separate L1 penalty to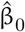.
2. To encourage DendroPRS to select a similar set of sparse effect sizes across all populations, we apply a group-lasso penalty to the set of parameters contributing to the effect size of each SNP, such that SNPs with a sparse effect size at the root are encouraged to remain sparse at the leaves.

Taken together with the prediction error of **y**, these three regularization penalties define our loss function ℒ.

Let β denote the set of all learnable parameters. Further, let ***μ***_*j*_ *∈* ℝ ^|*E*|+1^ denote the vector of parameters for SNP *j*, containing each of 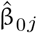 and 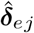 for all *e ∈ ε* :

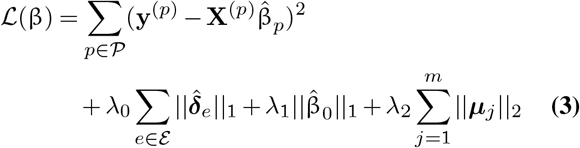

The three regularization terms are controlled by the hyperparameters *λ*_0_, *λ*_1_, *λ*_2_, which we tune via cross-validation. In the case of categorical phenotypes, an equivalent loss function can be defined by replacing the mean-squared-error (MSE) term with a cross-entropy term. While not considered in this current analysis, branch length annotations from *𝒯* can be easily incorporated by scaling *λ*_0_ inversely proportional to the length of each branch. We implement DendroPRS in PyTorch [Paszke et al., 2019] and optimize β via gradient-descent, using the Adam optimizer [Kingma and Ba, 2015]. As a future work, we could employ proximal gradient methods as in LASSO [Tibshirani, 1996] and Group-LASSO [Yuan and Lin, 2006] to get truly sparse estimates.

## Experimentation

The following protocols remain consistent across all experiments, unless specified otherwise. Hyper-parameters, including scaling factors of regularization terms and learning rate, are tuned via 5-fold cross-validation. Model evaluation is performed on a test set consisting of 30% of the samples, held-out from training and validation. For the quantitative phenotypes analyzed, the evaluation metric presented is the Pearson correlation coefficient between predictions ***ŷ*** and true ***y***.

The genotype, covariates, and target phenotypes are normalized to zero mean and unit variance. SNP selection criteria varies between experiments.

### Simulation Studies

We validated the ability of DendroPRS to fit population-specific effect sizes in a simulation setting. For the simulation procedure, we utilize actual genotype data from the 1000 Genomes Project (1KG) dataset. We *𝒫* define as the set of annotated “super” population groups: African (AFR, *n* = 661), South American (AMR, *n* = 347), East Asian (EAS, *n* = 504), European (EUR, *n* = 503), and South Asian (SAS, *n* = 489).

Human population history is still being actively elucidated with various tools and sources of evidence, from linguistics to global patterns of genetic variation [Cavalli-Sforza et al., 1988, Li et al., 2008, Duda and Zrzavý, 2016] and, more recently, with ancient DNA [Wohns et al., 2022]. While acknowledging that it is hard to represent this complex and interweaving history with a phylogenetic tree, this representation can still serve as a useful approximation of the global patterns found by previous studies. The tree used in our experimentation is displayed in Figure 1, and depicts the findings from Duda and Zrzavý [2016], with the simplifying assumption of constant branch lengths.

**Fig. 1.**
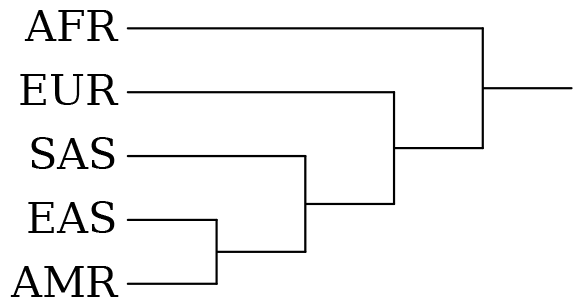
Phylogenetic tree with representative topology *𝒯* reflecting a global approximation of genetic and anthropological relationships between human continental populations. The leaves are labelled with the codes for the five “super” populations in the 1000 Genomes Project (1KG). Branch lengths are not to scale.

We used a set of 100 randomly chosen HapMap3 SNPs [Consortium et al., 2010] located on chromosome 22 with minor allele frequency (MAF) *>* 0.05. We simulated effect sizes using the magenpy package developed by Zabad et al. [2023]. Similar to the generative process outlined in the XPA method [Cai et al., 2021], effect sizes are drawn from a sparse mixture of multivariate Gaussian distributions, with covariance in effect sizes across populations scaled to be proportional to distance between the two corresponding nodes in the tree, while varying the minimum covariance in cross-population effect size (e.g. between the two most separated populations). For each set of simulated effect sizes, we simulated phenotypes exhibiting low, moderate and high levels of heritability, where the heritability level denotes the proportion of the simulated phenotype generated by the SNP features, the rest being Gaussian noise.

We compared the performance of DendroPRS against the baseline method of computing a single set of effect sizes across all populations via linear regression, with results summarized in Figure 2.

**Fig. 2.**
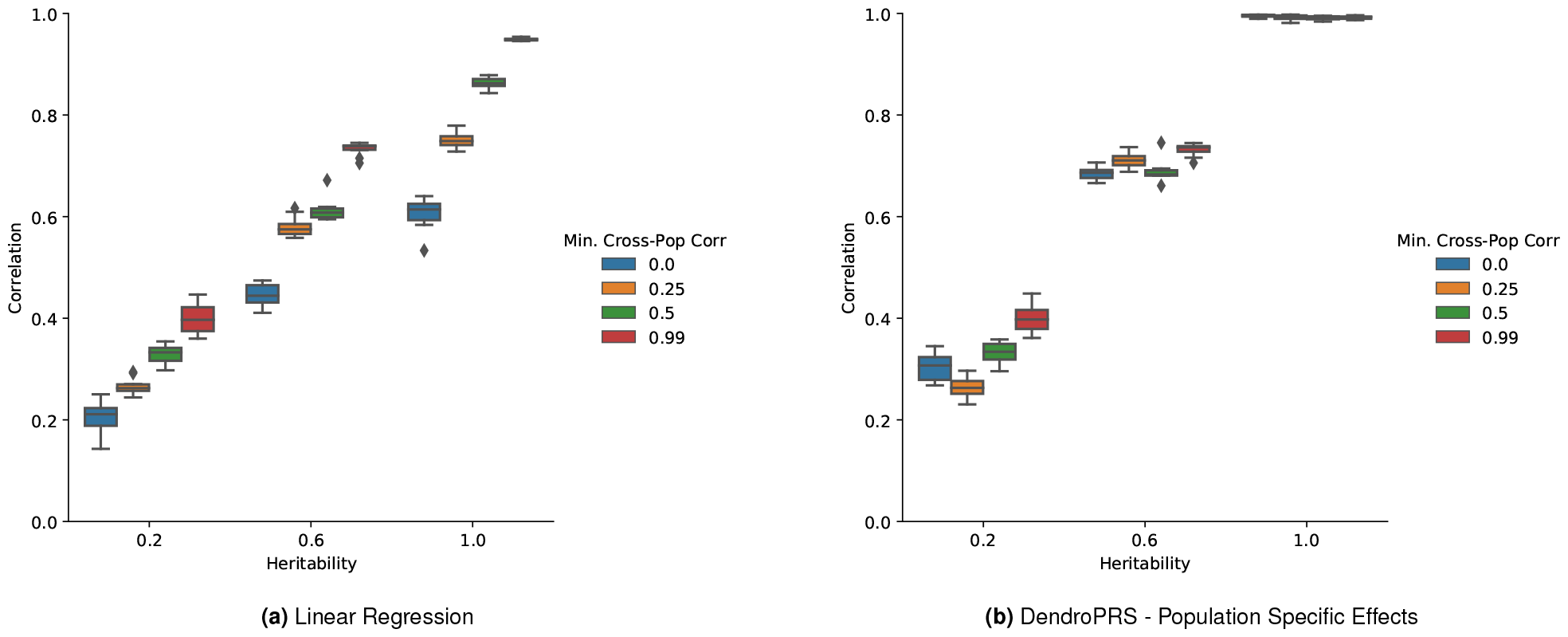
Performance of polygenic prediction approaches as a function of heritability and cross-population correlation in effect size. On the y-axis - Pearson correlation coefficient between simulated phenotypes and predictions made by fitting (left) a single set of effect sizes or (right) population-specific effect sizes with DendroPRS. On the x-axis, the heritability of the phenotype. Colour of bars indicate the minimum entry in the covariance matrix used for generating simulated effect sizes - i.e. the strength of correlation between the two least related populations.

For each of the simulated heritability settings, linear regression and DendroPRS perform similarly when the ground truth effect sizes are highly correlated across all populations. As the effect sizes become less strongly correlated across populations, the performance of the baseline method degrades rapidly, while DendroPRS remains stable, particularly in the high and moderate heritability settings. Taken together, these results provide evidence for DendroPRS’ ability to model population-specific effect sizes across different levels of heritability. Notably, in the setting where effect sizes are kept effectively constant across populations, DendroPRS exhibits no discernable disadvantage in comparison to the single effect-size model.

### Gene Expression Analysis

We next evaluated DendroPRS’s performance on real biological data. Our first study investigated the ability of DendroPRS to predict individual gene expression levels by estimating the regulatory effects of nearby SNPs, commonly referred to as expression quantitative trait loci (eQTL).

We extracted gene expression data from two ArrayExpress analyses of lymphoblastoid cells, from datasets containing individuals also included in the 1KG dataset. Gene expression measurements for African (*n* = 159), South American (*n* = 42), East Asian (*n* = 158) and South Asian (*n* = 75) individuals were extracted from the dataset compiled by Stranger et al. [2012]. Additionally, European samples (*n* = 358) were retrieved from the dataset presented by Lappalainen et al. [2013]. The two data sets were scaled, lognormalized, and combined. We focused our analysis on 649 genes whose expression were most highly correlated in a subset of 89 African individuals found in common between the two data sets. For those 89 African individuals, we retained only the expression data from Stranger et al. [2012], removing the duplicate samples from Lappalainen et al. [2013]. We used the same tree *𝒯* for the population groups as in Figure 1.

Genotype data was obtained from the 1KG dataset, focusing, for each gene, on the SNPs located within 50 kb of the transcription start site. We opted to use this distance cut-off, rather than the more standard 500 kb, as this captures the majority of SNPs likely to have a regulatory effect [Stranger et al., 2012], while filtering out many irrelevant SNPs which could have caused optimization issues for our methods, given the relatively small size of our dataset. Only SNPs with a MAF *>* 0.05 were retained.

For each of the 649 genes, we fit population-specific effect sizes using DendroPRS, and compared its performance in predicting gene expression levels against three baseline methods:

1. We fit population-specific effect sizes separately using only the subset of samples originating from each population (“Single-Pop.”).
2. We fit a single set of effect sizes using all populations jointly (“Pooled Pop.”).
3. We added a one-hot encoding of the population group to each sample, and fit a single set of effect sizes (“Pooled + Pop. Info”).

For each baseline method, L1-penalized regression is used to fit effect sizes. We evaluated each method’s average performance across 5 random train-validation-test splits of the data, with hyper-parameters tuned on the validation set, and performance evaluated on the test set.

For each method, we report the Pearson correlation between predicted gene expression and true expression levels, calculated separately within each population. Given that we expect the regulatory impact of SNPs to remain similar across populations in most genes, we are more interested in the subset of genes where there is a clear differentiation in performance, rather than the average performance across all genes. To identify cases where DendroPRS’s performance was clearly differentiated from a baseline method, we calculated 95% confidence interval (CI) for each correlation statistic. We used a Fisher Z-transformation [Fisher, 1937] to convert the observed correlations into approximately normally distributed estimated sample means, yielding the following formula where *r* is the Pearson correlation coefficient, *n* is the sample size and *c* = 1.96 is the critical value for 95% CI:

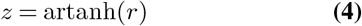

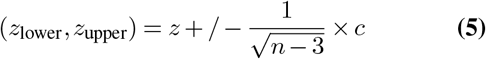

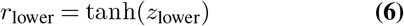

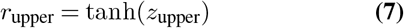

We recorded instances where the CI for DendroPRS and the comparison method had no overlap. Results are presented in Figure 3. DendroPRS broadly outperforms baseline method Single-Pop., exhibiting a total of 67 significantly improved genes across all populations, compared to only 15 genes where the Single-Pop approach was superior. Performance was improved in all populations, but particularly in the smaller non-European populations. This likely exhibits the limitations of fitting effect-sizes using population-specific data without access to sufficiently many samples, as can often be the case when analyzing groups that remain underrepresented in genetic datasets.

**Fig. 3.**
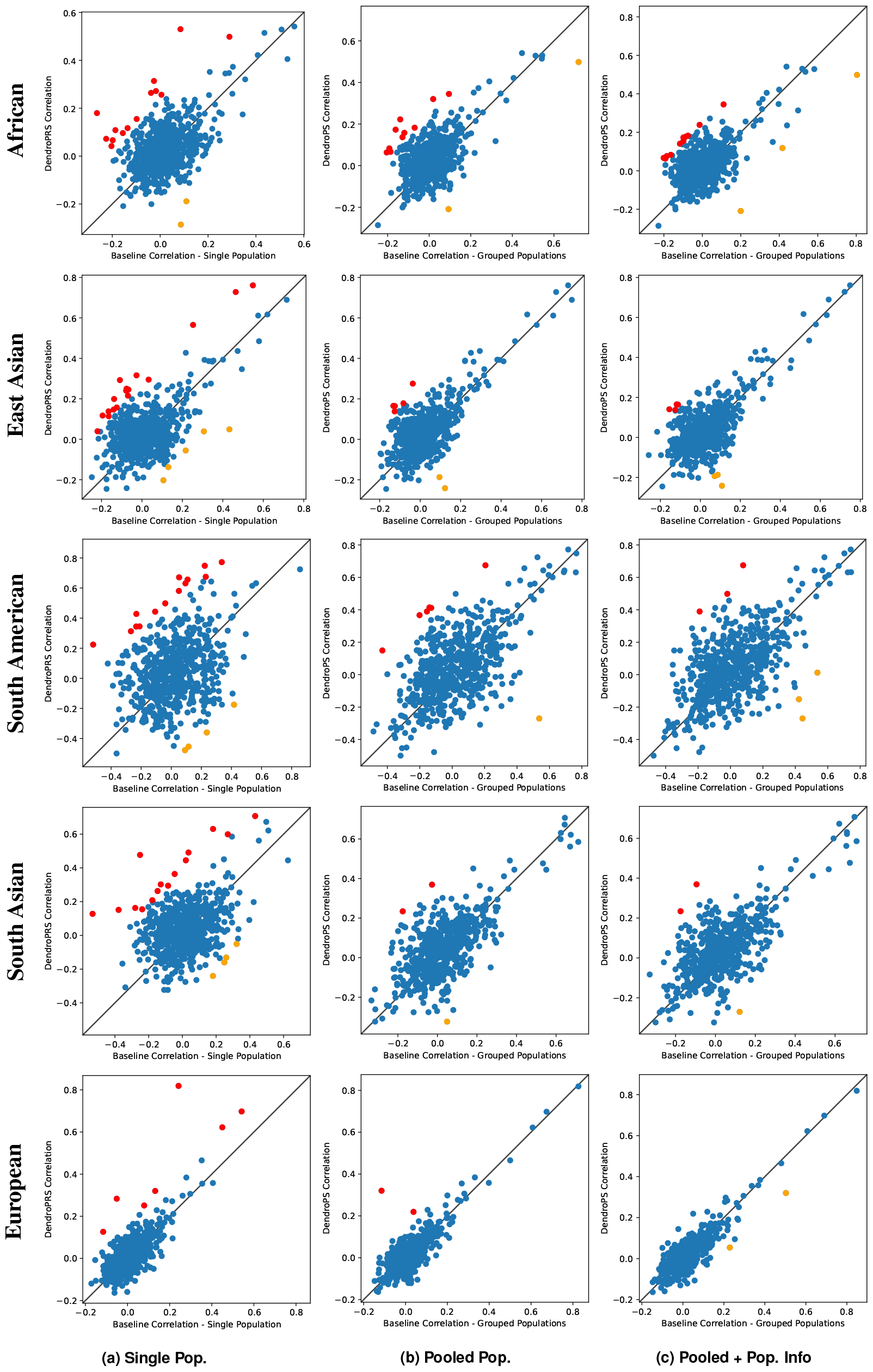
An analysis comparing the results of DendroPRS against 3 baseline methods, across 649 gene expression measurements within 5 populations. Genes marked in red are instances where the 0.95 CI of the correlation between true gene expression and DendroPRS predictions exceeds the comparison method within the respective population. Orange dots indicate instances of the inverse, where the baseline method is superior. Depicted in the first column, DendroPRS (y-axis) broadly outperforms the baseline method of computing training a separate prediction model for each population, suggesting that the reduced sample size when fitting single-population models is prohibitive. In comparison to training a single set of effect sizes across the pooled populations (middle column), DendroPRS exhibits notable advantages within the African and South American populations, and small advantages in other populations. The final column compares against the baseline method of fitting a model samples from all populations, each annotated with a one-hot encoding of population membership. DendroPRS exhibits significant advantages to this baseline method within the African population, and achieves approximately equivalent performance across other populations.

Similarly, DendroPRS out-performs baseline method Pooled Pop. in all populations, though by a tighter margin. There are a total of 25 genes across all populations where DendroPRS significantly improved expression prediction, in comparison to only 6 where the baseline method was superior. Most of the improvements exhibited by DendroPRS occurred within the African (10 improved genes), South American (6 improved genes) and East Asian (5 improved genes) populations. The limited number of instances where baseline method Pooled Pop. outperformed DendroPRS provides some assuring evidence that our phylogenetic regularization scheme is effectively preventing over-fitting even for the smaller populations.

Baseline method Pooled + Pop. Info was the strongest comparison method, having the ability to both leverage the full set of samples, and fit population specific terms via features capturing population information. DendroPRS modestly outperformed this method globally (20 improved genes to 12), but exhibited a notable benefit within the African population in particular, where it significantly improved prediction for 11 genes. This suggests that fitting African-specific intercept terms is not sufficient to accurately model gene expression, and that the ability afforded by DendroPRS to model changes to the regulatory effects of SNPs was additionally beneficial. Crucially, this analysis presents the opportunity to identify genes where there is particularly strong evidence of changes in regulatory architecture.

### Complex trait prediction from UK Biobank data

We next investigated the ability of DendroPRS to model the genetic effects underpinning highly polygenic traits across multiple populations at the biobank scale. The UK BioBank is the largest dataset readily available for PRS research, containing genome sequence data and extensive phenotype data from half a million individuals. Approximately 80% of the individuals are denoted as having “White British” ancestry, but many other ancestries are represented at the scale of several thousand individuals each, permitting the training and testing of cross-ancestry PRS models.

We studied six phenotypes: height, body mass index (BMI), high-density lipoprotein cholesterol levels (HDL), hip circumference, waist circumference, and forced expiratory volume (FEV1), all of which are both commonly studied and known to exhibit generalization difficulties in non-European populations [Duncan et al., 2019, Zabad et al., 2023].

The UKB collected self-declared ethnic background from all participants. The backgrounds are organized into a hierarchical dictionary that translates naturally into a tree structure, described in Data Field 21000 and Data Coding 1001 [Bycroft et al., 2018]. Self-declared ethnicity does not perfectly map to genetic ancestry, but it captures both genetic and environmental effects that may be relevant for modeling the genetics of complex traits. We opt for the simplest approach and directly translate the hierarchical data-encoding provided by UKB into a population tree for use with DendroPRS, acknowledging that this is an imperfect solution. In future work, a more sophisticated approach based on genetic clustering methods (e.g. [Prive, 2022, Diaz-Papkovich et al., 2023]) could yield further improvements.

We removed from consideration groups of mixed ancestry, as well as the Bangladeshi and “Other Black” populations due to their very small sample size. This yields a population tree with 9 leaves and three internal nodes, summarized in Table 1. We note that the ambiguous “Other Asian” population is grouped by UKB with South Asian populations under the “Asian” internal node, while the East Asian ancestry group, Chinese, is clustered separately.

**Table 1.** Summary of UKB populations included in our analyses and the corresponding number of samples [Bycroft et al., 2018]. Groupings into internal nodes defines the topology of *T* used to train DendroPRS.

| Population | Internal Node | Total Samples |
| --- | --- | --- |
| British | White | 472098 |
| Irish | White | 13992 |
| Other White | White | 17023 |
| Indian | Asian | 6108 |
| Pakistani | Asian | 1903 |
| Other Asian | Asian | 1853 |
| Caribbean | Black | 4606 |
| African | Black | 3460 |
| Chinese | Chinese | 1690 |

We selected SNPs relevant to each phenotype via the following procedure. First, universally across all phenotypes, we restricted analysis to HapMap 3 SNPs [Consortium et al., 2010] with MAF*>* 0.01. We then applied LD-pruning using the Plink2 indep-pairwise command with a window size of 100, step size of 5 and threshold of 0.2 [Chang et al., 2015]. For each phenotype, and separately for each super-group (White, Asian, Black, Chinese), we calculate p-values for SNP association via Plink2’s glm command [Chang et al., 2015] and retained SNPs that obtained a p-value *<* 5 *×*10^*−*8^ in at least one super-group. This enables the inclusion of SNPs that did not meet the significance threshold within the White British samples, but were significant within other populations. The number of SNPs selected for each phenotype is shown in Table 2.

**Table 2.** SNPs meeting inclusion criteria per UKB phenotype.

| Phenotype | Num. SNPs |
| --- | --- |
| Height | 608 |
| BMI | 243 |
| HDL | 48 |
| Hip Circumference | 237 |
| Waist Circumference | 290 |
| FEV1 | 74 |

In addition to SNP data, we include sex, age, and the first 10 Principal Components (PCs) of the genotype matrix as covariates in all analyses. Inclusion of the first 10 PCs in order to capture population structure is a standard practice in this type of analysis [Price et al., 2006], and their inclusion as covariates plays a similar role to the annotation of population membership in baseline method Pooled + Pop. Info in our eQTL experimentation. We note that the inclusion of sex and age as features in our model deviates slightly from the typical PRS workflow, in which these covariates are instead regressed out of phenotypes before fitting SNP effects. We opt to fit effects for covariates and SNPs simultaneously to enable DendroPRS to learn population-specific covariate effects, which may further improve predictive performance.

We set aside 30% of each ancestry group as a test set, and split the remaining samples into 70% training and 30% validation sets. We evaluate DendroPRS against two baseline methods:

- Lasso regression trained separately within each population (“Lasso - Single Pop”).
- Lasso regression trained on all data simultaneously (“Lasso - Pooled”).

Both baseline methods utilize the sklearn implementation of Lasso regression, optimized via coordinate descent [Pedregosa et al., 2011]. We note that in the case of single population Lasso, the baseline is still permitted to use SNPs that were identified using data from all populations.

All hyper-parameters are tuned on the validation set, including the alpha parameter for the Lasso regressors, the learning rate, *λ*_0_ and *λ*_1_ for DendroPRS. We fixed *λ*_2_ to 0 for all UKB experiments, due to negligible evidence of performance improvement throughout early experimentation. We trained DendroPRS for 300 epochs with an early-stopping criteria of 10 epochs, using mini-batches of size 128.

Figure 4 presents the Pearson correlation coefficient between predicted and true phenotype measurements demonstrated by DendroPRS and the baseline methods on the heldout test set. We assessed performance separately for each population within the study, with the final column in each sub-plot depicting the performance aggregated over all non-White populations.

**Fig. 4.**
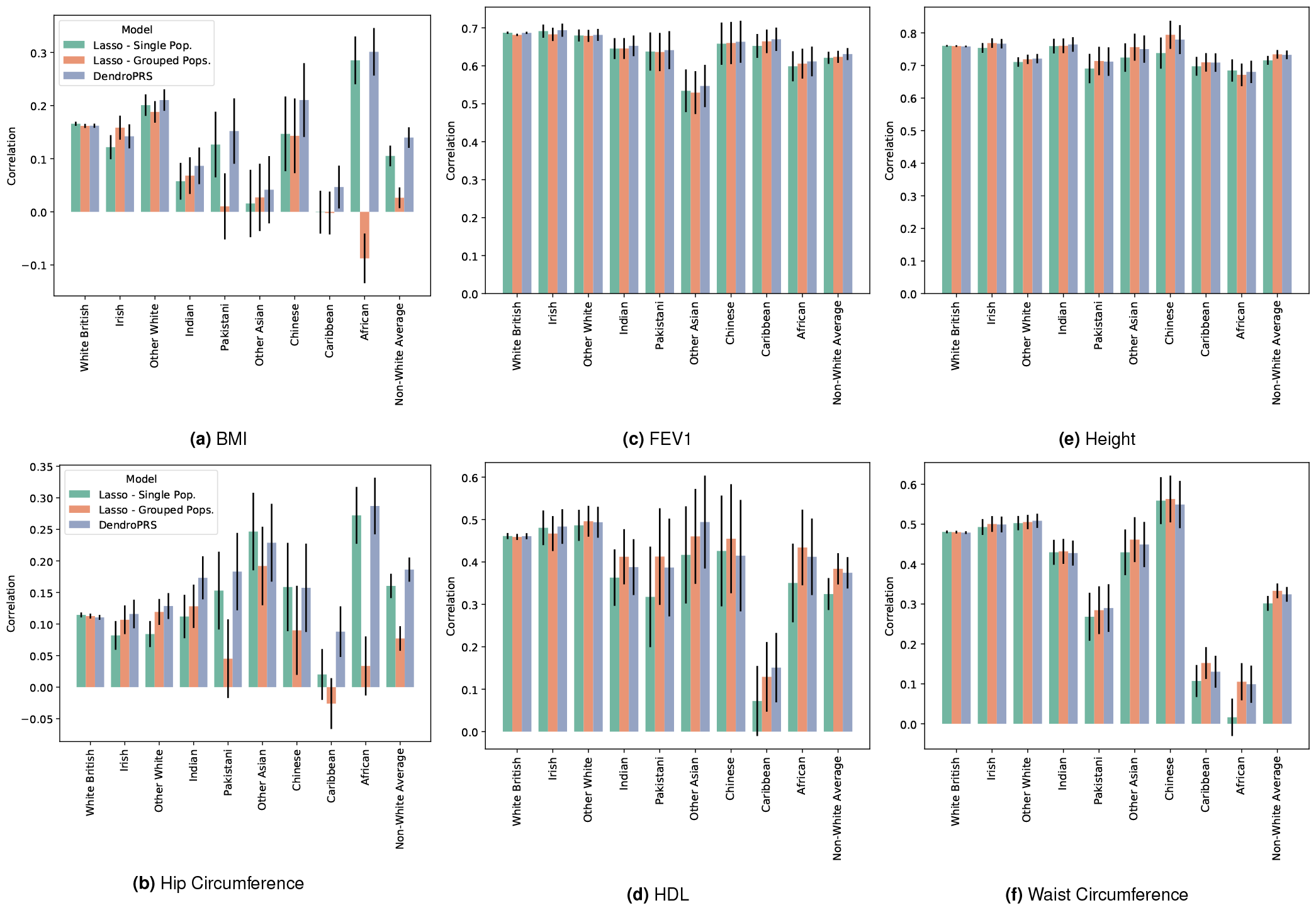
Performance of polygenic score methods on six quantitative phenotypes in the UK Biobank across different ethnic groups. The Pearson correlation coefficient between true and predicted phenotype is compared for DendroPRS and baseline methods (Lasso fit within a single population vs. grouped across all populations). The phenotypes analyzed are Body Mass Index (BMI), Hip Circumference, Forced Forced expiratory volume in 1 second (FEV1), High-density lipoprotein (HDL) cholesterol, Standing Height, and Waist Circumference. Error bars depict standard error of the Pearson correlation coefficient. DendroPRS exhibits broad benefits across non-White groups for the BMI and Hip Circumference traits, and additionally improves over single-population Lasso on the traits HDL and Waist Circumference. The strong results achieved by all methods on height and FEV1 are attributed in part to the inclusion of sex-differences within the phenotypes.

DendroPRS exhibits the most significant advantage on the tasks of predicting BMI and hip circumference. On BMI, DendroPRS improves correlation on aggregated non-White populations to 0.14 in comparison to 0.10 (Lasso - Single) and 0.03 (Lasso - Pooled). Improvements are particularly strong within Chinese, Pakistani, Caribbean, and African populations.

On hip circumference, DendroPRS attains a correlation of 0.19 on aggregated non-White populations, relative to 0.16 and 0.08 for Lasso - Single and Lasso - Pooled respectively. Gains are particularly strong within the Pakistani and Caribbean populations. DendroPRS also outperforms Lasso - Pooled within the African and Chinese populations.

For HDL, DendroPRS tracks the performance Lasso - Pooled quite closely. Improvements are obtained in comparison to Lasso - Single within non-White Average (0.32 improved to 0.37), particularly African, Caribbean, and Pakistani.

Similarly, DendroPRS outperforms Lasso - Single at the prediction of waist circumference within Africans; the three methods perform similarly within other populations.

All methods perform similarly on height and FEV1, with strong correlation levels exhibited across all populations. We attribute these elevated correlation levels to the inclusion of sex as a covariate within the PRS models, as both traits are known to have strong sex-specific differences in distribution.

### Variation in Effect Sizes Corresponds to Known Biological Mechanisms

The association between fat distribution and metabolic disturbance has been shown to be different in Europeans and Asians [Lindgren et al., 2009, Ma et al., 2019]. DendroPRS offers means to potentially identify the genetic causes of this difference. Our analysis of the BMI phenotype identifies rs11777976 as a SNP whose effect size changes significantly, both in sign and magnitude (top 1%), among populations. Whereas the major allele A is weakly positively associated with BMI in White and Black ancestry groups, DendroPRS assigns it a strong negative effect size within the Chinese population (*−* 0.07). This SNP is located in an intron of the MSRA gene [Kent et al., 2002], which has previously been associated with adipocity and BMI in various European populations [Ma et al., 2019]. Notably, the minor allele frequency at this SNPs is very high in Europeans (MAF close to 0.5), but very low in Asians (MAF= 0.04) [Sherry et al., 2001].

Additionally, DendroPRS assigns the covariate sex the strongest varying effect sizes of all features considered, with the direction of association flipped between the White and Chinese groups and other populations. This again aligns with previous findings of the relationship between sex and BMI varying within different populations[Karnes et al., 2021, Hendley et al., 2011, Stanford et al., 2019].

Together, these findings provide initial evidence that DendroPRS is capable of learning population specific effect sizes that accurately capture underlying sources of variation. They also demonstrate the potential future utility of DendroPRS’s approach for qualitative analysis of genetic variation between ancestry groups, potentially identifying fruitful research directions for future work.

## Conclusion

Robust polygenic risk scores prediction across diverse ancestry backgrounds is an ongoing challenge. In this work, we have presented DendroPRS, a novel tree-guided method for learning multi-ancestry polygenic risk scores. DendroPRS jointly learns effect sizes using data from all populations, leveraging phylogenetic-style regularization to prevent overfitting effect sizes on under-represented groups.

We investigated the performance of DendroPRS on three datasets: a simulation study, modelling eQTL effects on gene expression, and prediction of complex traits with UKB genetic data. In all cases, we found evidence of DendroPRS’ utility in fitting population-specific effect sizes. DendroPRS is able to identify numerous genes where expression can be predicted significantly more accurately by modelling population-specific regulatory effects. DendroPRS results on cross-ancestry eQTL may provide a valuable source of information for practitioners seeking to understand downstream phenotypes that vary across ancestry groups due to altered regulatory activity.

In the UKB analyses, we found that DendroPRS can outperform comparison methods across multiple traits and ancestry groups, with particularly strong gains attained for the Caribbean and African populations, demonstrating the potential value of tree-guided learning approaches for addressing generalization challenges within PRS.

Multiple opportunities exist for future work. DendroPRS should be scaled up to larger experiments involving many more SNPs, permitting optimal prediction of highly polygenic traits. This may require advances in regularization, such as branch or SNP specific penalty strengths, in order to avoid over-fitting on smaller populations. Our current implementation uses gradient descent to fit the effect sizes (Eq. 3). LASSO optimization schemes using coordinate ascent or proximal gradient methods could enable fitting truly sparse parameters, potentially improving both the ability to scale to more SNPs and interpretability of the estimated effect sizes. Finally, our framework could be extended to support admixed populations, by generalizing our tree-based implementation to support directed acyclic graphs, and thus allowing admixed populations to inherit effect size updates from multiple parent populations.

## Software and Data

An implementation of DendroPRS is available at https://github.com/BlanchetteLab/DendroPRS. Genotype and gene expression data for the eQTL analyses are publicly available on the 1KG website (https://www.internationalgenome.org/data). This research has been conducted using data from UK Biobank (Application 45551), a major biomedical database (https://www.ukbiobank.ac.uk).

## Author contributions statement

E.L. conceived of and implemented the method, and conducted experimentation. Y.L. and M.B. guided experimentation design and analysis. S.Z. assisted with simulation procedure and experiment design. All authors contributed to writing and editing the manuscript.

## Acknowledgments

We acknowledge the support of the Natural Sciences and Engineering Research Council of Canada (NSERC). Nous remercions le Conseil de recherches en sciences naturelles et en génie du Canada (CRSNG) de son soutien. The authors would like to thank Audrey Baguette, Yanlin Zhang, Dongjoon Lim, Ming Yang Zhou, Bowei Xiao, Dhanya Sridhar, Jason Hartford, Alex Tong, Mathieu Bourgey and Zichao Yan for helpful conversations.

